# Metatranscriptomics supports mechanism for biocathode electroautotrophy by “*Candidatus* Tenderia electrophaga”

**DOI:** 10.1101/074997

**Authors:** Brian J. Eddie, Zheng Wang, W. Judson Hervey, Dagmar H. Leary, Anthony P. Malanoski, Leonard M. Tender, Baochuan Lin, Sarah M. Strycharz-Glaven

## Abstract

Biocathodes provide a stable electron source to drive reduction reactions in electrotrophic microbial electrochemical systems. Electroautotrophic biocathode communities may be more robust than monocultures in environmentally relevant settings, but some members are not easily cultivated outside of the electrode environment. We previously used metagenomics and metaproteomics to propose a pathway for coupling extracellular electron transfer (EET) to carbon fixation in “*Candidatus* Tenderia electrophaga”, an uncultivated but dominant member of the Biocathode-MCL electroautotrophic community. Here we validate and refine this proposed pathway using differential metatranscriptomics of replicate MCL reactors poised at the growth potential 310 mV and the suboptimal 470 mV (vs. standard hydrogen electrode). At both potentials, transcripts from “*Ca*. Tenderia electrophaga” were more abundant than from any other organism and its relative activity was positively correlated with current. Several genes encoding key components of the proposed “*Ca*. Tenderia electrophaga” EET pathway were more highly expressed at 470 mV, consistent with a need for cells to acquire more electrons to obtain the same amount of energy as at 310 mV. These included *cyc2*, encoding a homolog of a protein known to be involved in iron oxidation, confirmed to be differentially expressed by droplet digital PCR of independent biological replicates. Average expression of all CO_2_ fixation related genes is 1.23-fold higher at 310 mV, indicating that reduced energy availability at 470 mV decreased CO_2_ fixation. Our results substantiate the claim that “*Ca*. Tenderia electrophaga” is the key MCL electroautotroph, which will help guide further development of this community for microbial electrosynthesis.

**IMPORTANCE:** Bacteria that directly use electrodes as metabolic electron donors (biocathodes) have been proposed for applications ranging from microbial electrosynthesis to advanced bioelectronics for cellular communication with machines. However, just as we understand very little about oxidation of analogous natural insoluble electron donors, such as iron oxide, the organisms and extracellular electron transfer (EET) pathways underlying the electrode-cell direct electron transfer processes are almost completely unknown. Biocathodes are a stable biofilm cultivation platform to interrogate both the rate and mechanism of EET using electrochemistry and study the electroautotrophic organisms that catalyze these reactions. Here we provide new evidence supporting the hypothesis that the uncultured bacterium “*Candidatus* Tenderia electrophaga” directly couples extracellular electron transfer to CO_2_ fixation. Our results provide insight into developing biocathode technology, such as microbial electrosynthesis, as well as advancing our understanding of chemolithoautotrophy.

## INTRODUCTION

Biocathodes are bioelectrochemical systems (BES) in which microbial electrode catalysts use the electrode as an electron donor to drive cellular metabolism. Over the last decade biocathodes have been explored for improving energy recovery in microbial fuel cells (MFCs), electrode driven bioremediation, and more recently to produce chemicals in a process known as microbial electrosynthesis (reviewed in (1, 2)). Despite widespread interest in biocathodes for biotechnology applications, little is understood about the underlying mechanisms of extracellular electron transfer (EET) for biocathode microorganisms. The *Marinobacter*-*Chromatiaceae*-*Labrenzia* (MCL) biocathode is a self-regenerating, self-sustaining, aerobic microbial community and has served in our lab as a model system for the exploration of electroautotrophic microbial communities using an omics approach (3, 4). MCL forms a heterogeneously distributed biofilm with clumps of up to 20 µm thick (4) with reproducible electrochemical features following inoculation of a portion of the biofilm into a new reactor. It is proposed that MCL, which operates under aerobic conditions, reduces O_2_ with electrons supplied solely by the cathode, directing a portion of the acquired energy and electrons for autotrophy. Cyclic voltammetry (CV) reveals a sigmoid-shaped dependency for O_2_ reduction catalytic (turnover) current on electrode potential (5). It is proposed that this dependency reflects Nernstian behavior of the heterogeneous electron transfer reaction (across the biofilm/electrode interface) that is mediated by a redox cofactor, which is fast, reversible and not the rate limiting step in electron uptake form the cathode by the biofilm (4, 6, 7). Electroautotrophic growth is assumed for MCL based on an increase in biomass correlated to increasing current, a lack of organic carbon in the bioelectrochemical reactor, and identification of an active Calvin-Benson-Bassham cycle (CBB) (8). Electroautotrophic growth of isolates at potentials > −100 mV vs. SHE has thus far only been demonstrated for *Mariprofundus ferrooxydans* PV-1 and *Acidithiobacillus ferrooxidans* (9, 10), while other autotrophs require supplemental energy from light or hydrogen for initial growth on a cathode (11, 12). However, recent reports indicate that communities containing *Gammaproteobacteria* using a high potential biocathode as an energy source can be reproducibly enriched (13, 14).

Previous metagenomic and proteomics studies of MCL have led to the identification of putative EET and CO_2_ fixation mechanisms (8, 15) although efforts at cultivation have not yielded the proposed electroautroph, “*Candidatus* Tenderia electrophaga” (16). Other work has resulted in isolates from biocathode enrichments, but these have not been autotrophic (17, 18). Metaproteomic analysis suggests that proteins that may be involved in EET are expressed at high levels in the biofilm, including a homolog of Cyc2, a protein known to be involved in Fe^+2^ oxidation in *A. ferrooxidans* (8). Subsequent metaproteomic analysis of the biofilm at two different electrode potentials showed that some components of the electron transport chain (ETC) are differentially expressed including an ortholog of Cyc1, thought to be involved in Fe^+2^ oxidation in *M. ferrooxydans* PV-1 (19), and a hypothetical protein homologous to the terminal oxidase cytochrome cbb_3_ subunit CcoO (15). More precise quantification of the changes in gene expression obtained using RNA-seq to quantify the relative abundance of transcripts (20) can be used to understand the molecular mechanisms used for growth on the cathode.

Building upon our previous work (4, 8, 15), we applied RNA-seq to MCL to assess gene expression of proposed EET and CO_2_ fixation pathways for “*Ca*. Tenderia electrophaga” when the applied electrode potential is adjusted from 310 mV vs. SHE (optimal for growth) to 470 mV (suboptimal for growth). This 160 mV increase in potential decreases the ΔG^0'^ for the reduction of O_2_ by about one third, and from previous metaproteomic experiments run under identical conditions is expected to result in changes in gene expression to compensate for the change in energy availability. These results support previously hypothesized roles of some protein complexes (8, 15, 19), and reveal possible EET roles for other proteins. We also provide further evidence that “*Ca*. Tenderia electrophaga” is the keystone species in this community and is strongly associated with current production. These results provide further understanding of the molecular mechanisms involved in electroautotrophic growth on a cathode, enabling the use of members of this community to develop biocathodes for applications including synthesis of value added compounds or biofuels as well as potential ET components for microbial bioelectronics.

## RESULTS

### Biocathode-MCL Metatranscriptome

Eight BES were grown under previously described standard conditions at an applied electrode potential of 310 mV SHE. Seven of the eight biological replicates reached maximum current production within five days (Table 1, Fig. S1). Replicate S2A took two days longer. The midpoint potential (E_M_) of each BES measured by cyclic voltammetry (CV) once steady-state current was achieved was identical to that previously reported (4, 8, 15) at ca. 440 mV vs. SHE (Table 1). Following CV, the electrode potential was either returned to 310 mV or adjusted to 470 mV, a potential at which we expect the biofilm to recover less energy per electron based on previous experiments.

**Table 1.**
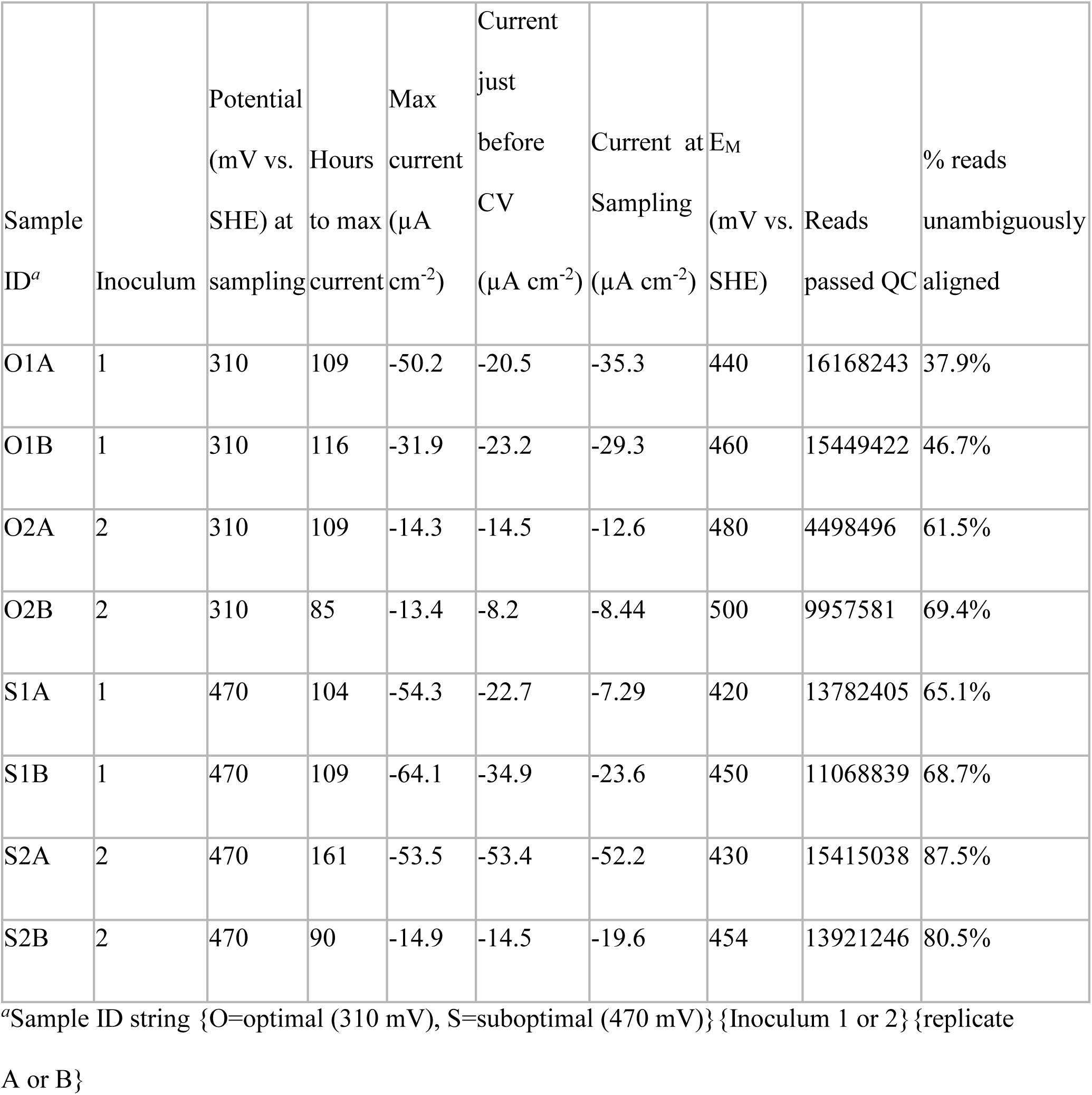
Sample and sequencing summary.

MiSeq mRNA sequencing yielded between 4.5 – 16.2 million reads per replicate sample that passed quality control steps and were used for read alignment to the metagenome, of which 37.9-87.5% could be unambiguously aligned (Table 1). The relative activity of MCL constituents was assessed as the proportion of reads mapping to a metagenome bin normalized by relative length of bin. Using this metric, MCL constituents previously implicated in key roles in the biocathode – “*Ca*. Tenderia electrophaga”, *Marinobacter* sp. strain CP1, and *Labrenzia* sp. strain CP4. – were highly active in all samples at both potentials. Ten other metagenome bins were active in all eight samples using a cutoff of 0.01% of relative activity. Two of these, *Parvibaculum* sp. and *Kordiimonas* sp. had activity similar to that of *Labrenzia* sp. and *Marinobacter* sp., indicating that they may also have significant roles in the biocathode community. “*Ca*. Tenderia electrophaga” was the most active organism in all eight samples, comprising 53% to 84.3% of activity (Fig. 1). *Marinobacter* sp., *Labrenzia* sp., *Kordiimonas* sp., and *Parvibaculum* sp. average 24.6% of the remaining activity. These five organisms make up the core of the biofilm community, on average accounting for 93.6% of community activity. Two-tailed Student's T-tests indicated that no organism has a significant change in activity between the two potentials tested (p > 0.09 for all bins).

**Figure 1.**
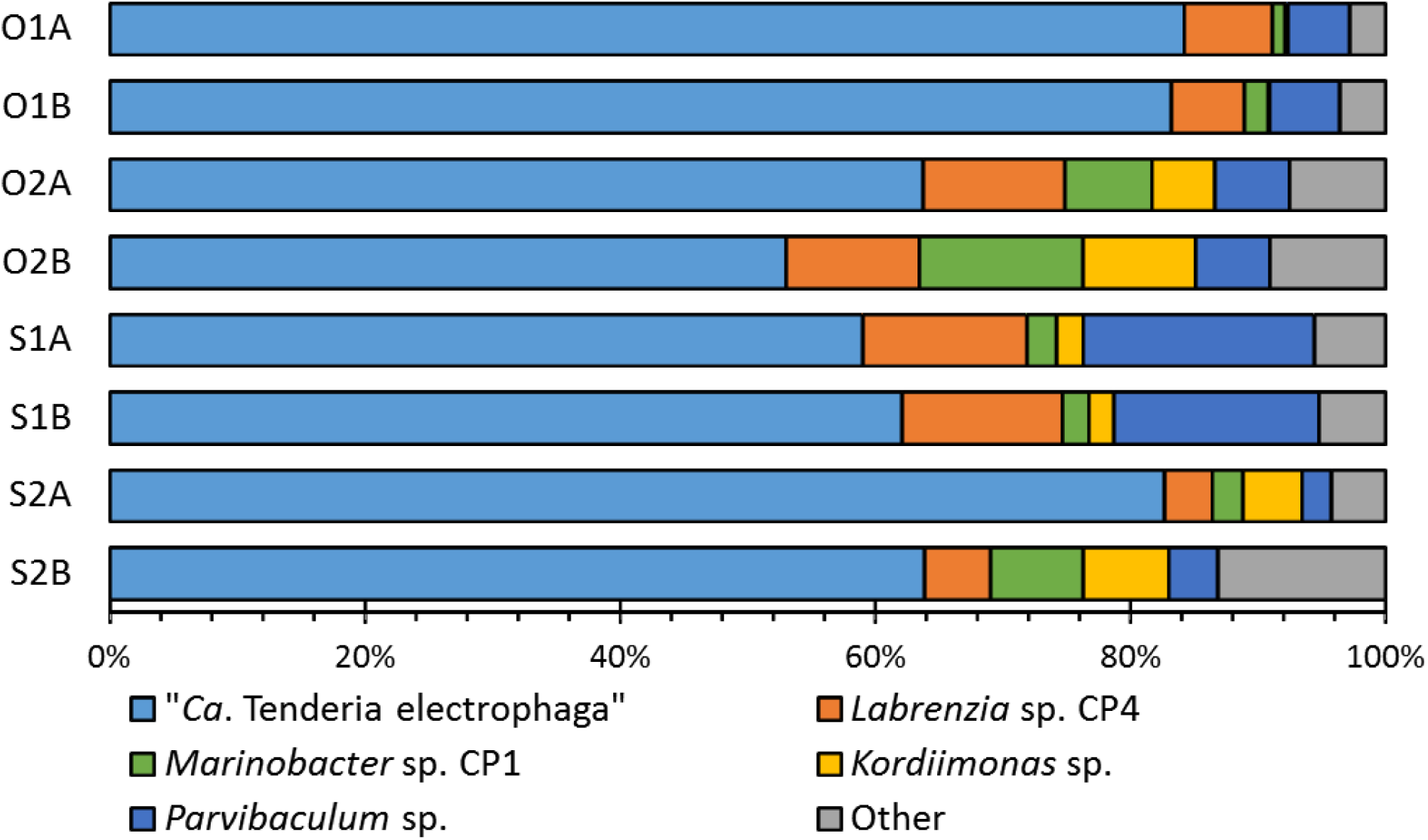
Relative activity of the five major constituents of Biocathode-MCL. Proportion of reads matching a genome or bin normalized by the length of the genome or bin was taken as a proxy for activity. Sample labels are as in Table 1.

### Correlation of “Ca. Tenderia electrophaga” transcriptional activity to current density

Activity of “*Ca*. Tenderia electrophaga” was strongly correlated to current density at both 310 mV and 470 mV (Fig. 2A) with Pearson correlation coefficients of 0.975 and 0.967 respectively. Activity of the four most active heterotrophic organisms, *Labrenzia* sp., *Marinobacter* sp., *Parvibaculum* sp., and *Kordiimonas* sp. was negatively correlated, or not correlated with current density (Fig. 2B-D). Previous metagenomic and metaproteomic data indicated that “*Ca*. Tenderia electrophaga” is likely an autotrophic organism capable of performing EET (8, 15) and the correlation of current density to activity supports this theory. For this reason, we concentrated our analysis of metatranscriptomics data on “*Ca*. Tenderia electrophaga” to focus specifically on how direct EET is linked to CO_2_ fixation.

**Figure 2.**
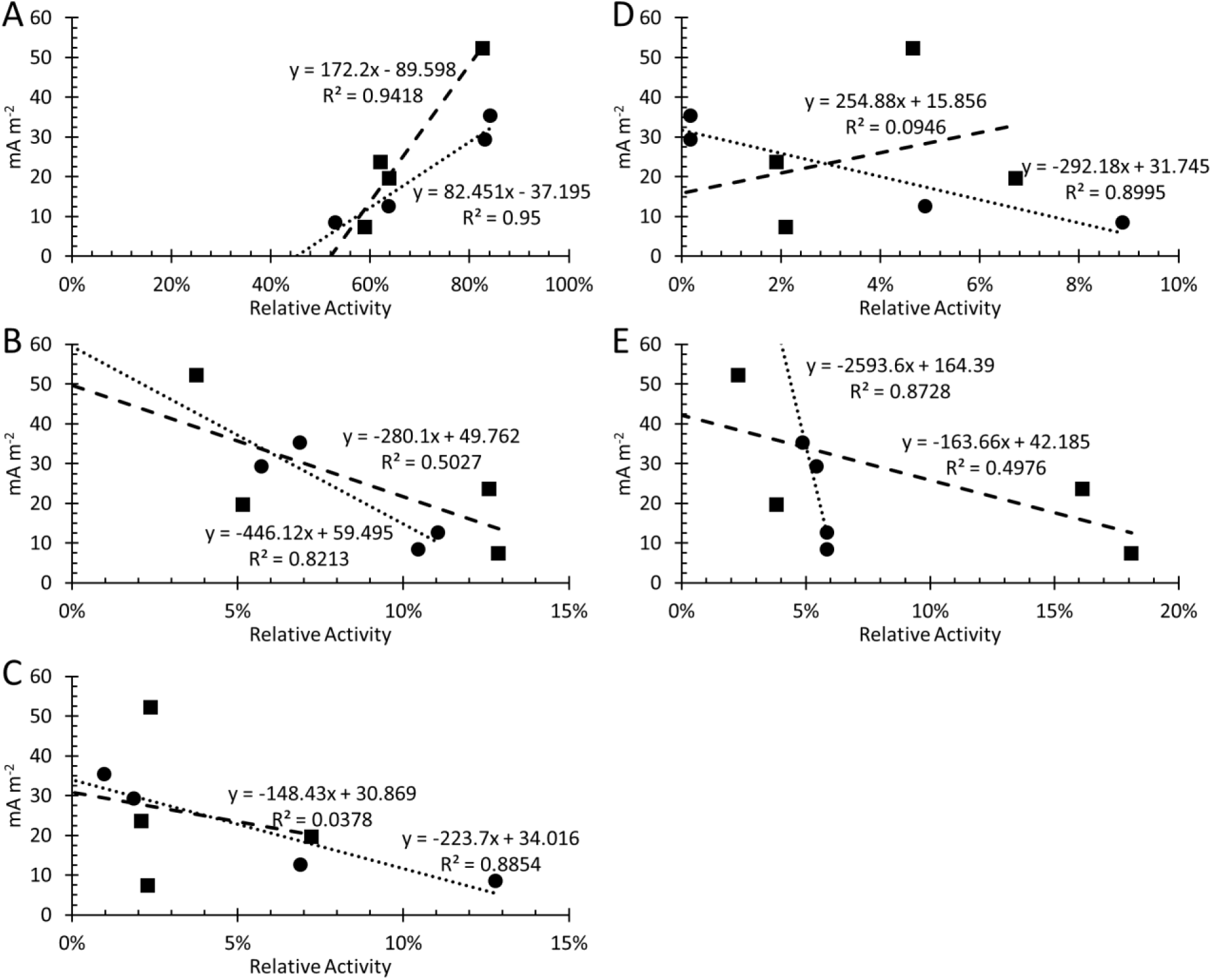
Activity of top five Biocathode-MCL constituents vs. current density at the optimal and suboptimal electrodes potentials. *“Ca*. Tenderia electrophaga” (A) has positive correlations with increasing negative current at both the optimal (circles) and suboptimal (squares) potentials. Other active members of the community: *Labrenzia* sp. CP4 (B), *Marinobacter* sp. CP1 (C), *Kordiimonas* sp. (D), and *Parvibaculum* sp. (E) had negative or insignificant correlation at both potentials. Linear regression lines are shown, along with the equation of the lines and R^2^ values.

### Different Pathways for Electron Transport Are Used at Different Potentials

Previous metagenomic and metaproteomic analysis of Biocathode-MCL revealed proteins for carbon fixation through the CBB cycle and a putative EET pathway in “*Ca*. Tenderia electrophaga” (8). Statistical analysis of protein expression revealed that nine proteins in “*Ca*. Tenderia electrophaga” were detected more often at one potential versus another, but methodological limitations meant that this was likely to be a small fraction of the response to changing potential (15). Efforts to cultivate a representative isolate of “*Ca*. Tenderia electrophaga” from MCL have thus far been unsuccessful, suggesting that other community constituents satisfy unknown requirements for its growth. We therefore used metatranscriptomics to obtain higher resolution analysis of differentially expressed mRNA levels for proteins suspected to be involved in electroautotrophic growth. This allowed us to test the hypothesis that expression of protein coding genes involved in respiration and energy flux in the primary electroautotroph is directly influenced by the potential of electrons supplied by the cathode. A complete list of genes we propose to be associated with these pathways is presented in Table S1.

Electrochemical measurements indicate redox dependent direct electron transfer (DET) is occurring between the electrode and “*Ca*. Tenderia electrophaga” (4, 5), therefore, we searched the metatranscriptome for evidence of an EET conduit whose expression may be affected by the change in electrode potential. The *cyc*2 homolog in “*Ca*. Tenderia electrophaga” (Tel_03480) was predicted to have a role in EET (8) based on its proposed involvement in iron oxidation and was significantly differentially expressed, with relative transcript levels 1.8-fold higher at 470 mV. The gene for a predicted hexaheme lipoprotein (Tel_04230) was significantly more highly expressed at 470 mV, raising the possibility that this protein is involved in electron transfer. An undecaheme *c*-cyt (Tel_16545) was previously identified as a possible route for EET in “*Ca*. Tenderia electrophaga” due to the large number of predicted heme binding sites, known to be important for EET in *Shewanella* and *Geobacter spp*., and its conservation among other EET capable organisms (8). The gene for this protein was not differentially expressed.

It is thought that soluble, periplasmic cytochromes mediate ET between the outer membrane and the cytoplasmic membrane bound ETC (21, 22). Several genes previously noted from “*Ca*. Tenderia electrophaga” which may encode proteins involved in transferring electrons across the periplasm to ETC were examined for changes in expression between the two electrode potentials. Expression was higher for three tri-heme cytochromes (Tel_16515, Tel_16520, Tel_16530) at 310 mV, but was not statistically significant (p > 0.01). Overall, these genes are among the most highly expressed in “*Ca*. Tenderia electrophaga”, indicating their importance for growth at the cathode. They also appear to be co-transcribed with a gene encoding a tetratricopeptide repeat (Tel_16525), which are known to be involved in protein:protein interaction and may be involved in aligning cytochromes in a bridge configuration as proposed for other EET capable bacteria (23). Peptides from this tetratricopeptide repeat protein were previously found to be significantly more abundant at the suboptimal potential (15), although in the metatranscriptomic data, this cluster of genes is slightly more highly transcribed at 310 mV.

A di-heme cytochrome c_4_, Cyc1, has been proposed to be a periplasmic electron shuttle in the iron oxidizing bacterium *M. ferrooxydans* (19). The gene encoding this protein is found in “*Ca*. Tenderia electrophaga”, in a region displaying synteny with two contigs from the *M. ferrooxydans* genome (Figure S2). Like the tri-heme cytochromes, it is very highly expressed at both potentials, but is not significantly differentially expressed. Other genes for potential periplasmic electron carriers identified from the metatranscriptome that were not previously implicated in MCL EET included a monoheme *c*-cyt (Tel_12755). This gene has the largest change in expression of any annotated cytochrome in “*Ca*. Tenderia electrophaga” (3-fold more highly expressed at 310 mV) suggesting that it may be important for electron transfer at this lower potential. It is homologous to genes for proteins in several iron oxidizing bacteria, including *Sideroxydans* (45% amino acid identity), *Gallionella* (42% identity), and *Acidithiobacillus* (42% identity) species. This may make it an important gene to consider in models of iron oxidation.

Regardless of the pathway used, e^−^ need to reach the cytoplasmic membrane bound ETC, which is the main mechanism for conservation of energy via chemiosmotic gradient formation (24). Metatranscriptomic analysis of the ETC components does not indicate large changes in the expression of these genes, which suggests that the same ETC components are used at both electrode potentials. Small (<1.3 fold) changes in the ratios of expression for whole pathways may represent subtle adjustments in the ratio of electrons going down each path (Table S1). Whether this is in addition to or instead of the pathway being selected by potential of the electron donor in a manner reminiscent of recent work on electron acceptors in *Geobacter sulfurreducens* remains to be determined (25, 26).

Most genes encoding components of the ETC were not significantly differentially expressed, although many had smaller but consistent changes in expression between the two potentials. For example, the genes encoding the Alternative Complex III (ACIII), the NADH:ubiquinone oxidoreductase complex (NUOR), and a homolog of Cyc1 from *M. ferrooxydans*, thought to be responsible for transferring electrons from the outer membrane to the cytochrome cbb_3_ oxidase (19), were not differentially expressed. However, hierarchical clustering based upon the slight changes of expression among these complexes (Fig. 3) suggests that they are functionally linked. This supports a role as the reverse electron transport (RET) chain, linking EET to the reduction of NAD(P)^+^ for fixation.

**Fig. 3.**
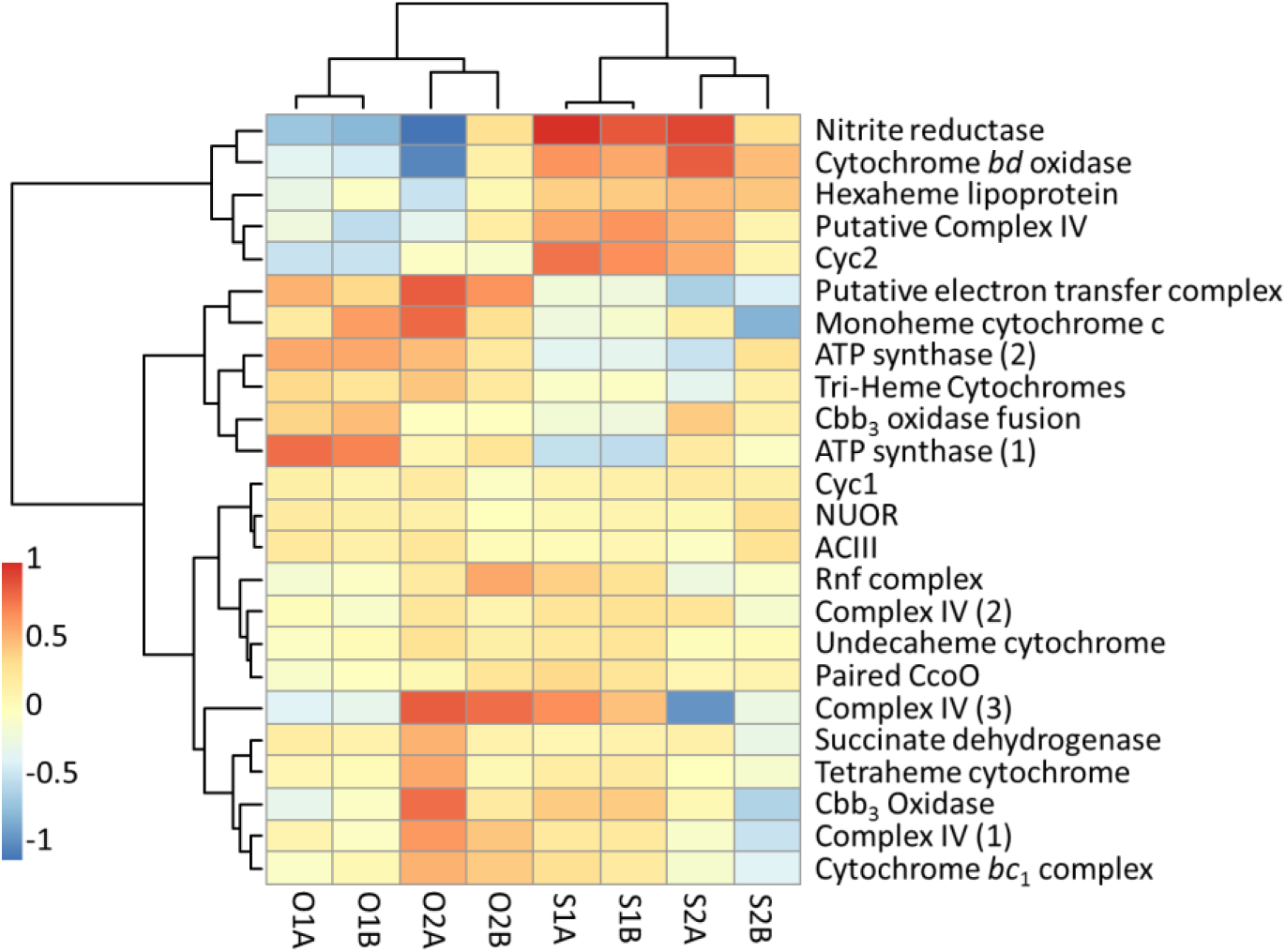
Hierarchical clustering of EET and ETC components by change in expression. Expression is normalized to 0 between biological replicates to focus on change in expression between 310 mV and 470 mV. Details about genes in each complex are found in Table S1. Sample identifiers are as in Table 1.

Only one operon in “*Ca*. Tenderia electrophaga” contains all four of the canonical genes for a bacterial cytochrome cbb_3_ oxidase (*ccoNOPQ*), but it has relatively low expression. However, the genome also encodes several proteins homologous to terminal oxidases (Complex IV) that could potentially reduce O_2_ and generate proton motive force (PMF). Of these, the genes encoding Complex IV-2 had some of the highest total expression of any genes, and clustered with the undecaheme cytochrome and two additional copies of *ccoO* (Fig. 3) which suggests that this represents a way for electrons to enter the cell and be used to reduce O_2_.

Among genes with a large change in expression, *cyc2* clustered with a gene cluster encoding a possible nitrate reductase and a complex including a *ccoO* homolog and a copper containing plastocyanin homolog labeled as Putative Complex IV. However, this cluster of genes lacks a protein that could function as an analog of the membrane associated proton pumping CcoN. Thus it is unlikely to make a substantial contribution to PMF. All of the genes encoding components of the two ATP synthase operons were more highly expressed at 310 mV, many with a false discovery rate (FDR) of <0.01, indicating that energy levels in “*Ca*. Tenderia electrophaga” are dependent upon the electrode potential.

Expression of 19 genes was quantified using droplet digital-PCR (ddPCR) to verify differential expression of several genes hypothesized to be important for growth on the cathode (Table S2). These measurements we performed on an independent set of eight MCL biocathodes, approximately one year after the original reactors. Only *cyc2* was found to be differentially expressed in the both ddPCR results and the metatranscriptome. Variability between inocula and the lack of an adequate method for normalizing to account for differences in the biofilm community may have reduced the sensitivity of the assay.

### Effect of potential on carbon fixation and central carbon metabolism

The “*Ca*. Tenderia electrophaga” genome contains all the genes necessary for CO_2_ fixation via the CBB cycle, including two forms of Rubisco – one associated with carboxysomes (Form IAc) and one not associated with carboxysomes (Form IAq) (8). The lower energy yield per electron from a cathode poised at 470 mV would result in less energy available for CO_2_ fixation, so the central carbon metabolism of “*Ca*. Tenderia electrophaga” was examined for changes in expression. Genes which could be involved in more than one pathway, for example, CO_2_ fixation and the pentose phosphate pathway, were separated where possible by examining co-localization and changes in expression between the eight samples. Genes with ambiguous functional roles that could not be assigned to a single pathway were excluded from this analysis. Most components of central carbon metabolism, including the CBB cycle were more highly expressed at 310 mV, although the absolute magnitude of the change was relatively small (Fig. 4, Table S3). Only four genes were differentially expressed, ribose 5-phosphate isomerase A and ribulose-phosphate 3-epimerase which catalyze anaplerotic reactions in the CBB cycle, and 6-phosphogluconate dehydrogenase and glucose-6-phosphate 1-dehydrogenase which catalyze components of the pentose phosphate pathway that supply metabolic precursors for biosynthesis. However, when examining all genes in a given pathway, they were mostly consistent, indicating that the pathways are likely genuinely more highly expressed. The exceptions are the form IAc Rubisco genes and carboxysome genes. These were more highly expressed at 470 mV, which suggests a switch to a more efficient enzyme under energy limitation. The whole CBB CO_2_ fixation cycle was 1.23-fold more highly expressed at 310 mV.

**Fig. 4.**
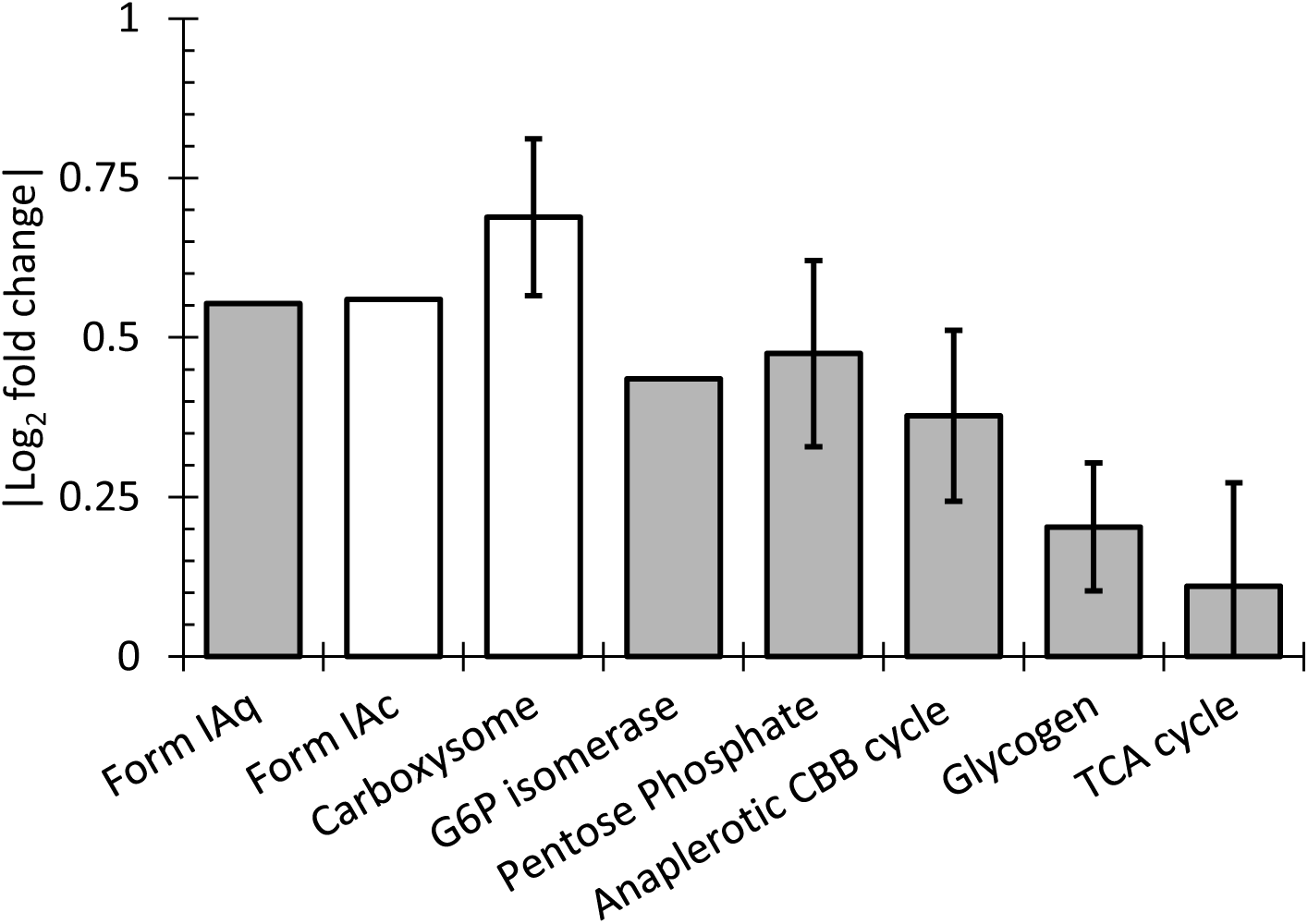
Change in expression of central carbon metabolism genes. Open boxes are more highly expressed at 470 mV, gray boxes are more highly expressed at 310 mV. Error bars indicate standard deviation of log2 fold change of pathways containing three or more genes. Details of the genes included in each category, and their expression can be found in Table S3.

## DISCUSSION

Our characterization of the Biocathode-MCL community by metatranscriptomics agrees with previous metaproteomic and 16S rRNA amplicon characterization of Biocathode-MCL that there is substantial variability in the abundance of specific bacteria, even between seemingly identical reactors inoculated on the same day with the same inoculum (15). However, as in the previous studies, changing the electrode potential from 310 mV to 470 mV for 52 hours after the community had developed did not result in significant changes in the average relative abundance of biocathode constituents between the two potentials. While the inherent variability in relative abundance between replicate reactors makes analysis of the transcriptome data challenging, differences that we found are more likely to be robust because they are visible above the noise of such large biological variability when using paired, replicated samples as in Leary et al. (15).

The magnitude of changes in gene expression were relatively small compared to other transcriptomic studies (e.g. (27, 28)). However, in one of those studies, replicate samples were acquired by repeatedly sampling the same biofilm, which may have inflated the reported statistical confidence, and in the other no replicates were sampled. The 52 hours adjustment period after changing the potential likely decreased the magnitude of the transcriptional response, but it was chosen to facilitate comparison to proteomics data (15). On a shorter timescale, transcription of genes for upregulated pathways may experience a burst, as the cells adjust to the new regulatory regime, but would relax back to the new steady state once the proper balance of proteins has been reached. Also, the fundamental metabolism of “*Ca*. Tenderia electrophaga” does not change. Electrons are still being transferred from the cathode to the ETC, and CO_2_ fixation is still the sole carbon source.

Our results suggest that “*Ca*. Tenderia electrophaga” is primarily responsible for EET linked to CO_2_ fixation in Biocathode-MCL biofilms. The dominance of “*Ca*. Tenderia electrophaga” transcriptional activity and its correlation to current density suggests that it can account for the majority of current. Further substantiating this hypothesis is the recent repeated identification of unclassified bacteria, thought to be within the *Chromatiales,* which were dominant members of freshwater biocathode communities (13, 14). While *Marinobacter* spp. are capable of cathode oxidation (17, 18, 29), isolates of *Marinobacter* sp. and *Labrenzia* sp. from the MCL-community are only capable of weak current production at the biocathode or iron oxidation in monoculture (8).

Based on a prior analysis of Biocathode-MCL using low scan rate CV, we predicted current to be approximately halved when the electrode potential was switched from 310 mV to 470 mV (15). However, as previously observed (15), for biocathode-MCL, switching to and maintaining a more positive potential (470 mV) over a much longer time period (>50 h) resulted in an increase in current attributed to O_2_ reduction after the initial adjustment period. This increase in current is interpreted as the result of the need to make up for the decrease in energy available per electron at the higher potential. Using a derivation of the Nernst Equation (ΔG = nFΔE^0'^) to estimate the theoretical yield of O_2_ reduction to H_2_O using an electron donor at a potential of 310 mV, yields −47 kJ/mol e^−^, requiring an additional ~61 kJ/mol e^−^ to reduce NAD(P)^+^. Assuming that the energy required to pump protons across the membrane is ~21 kJ/mol, at the optimal potential “*Ca*. Tenderia electrophaga” could export two H^+^ per e^−^, and would need to use three H^+^ to reduce NAD(P)^+^. This results in a theoretical balance for electron utilization by the forward vs. reverse electron transport pathways of ~ 60%/40% to produce NAD(P)H. More electrons would need to go to the forward path to generate a proton gradient for ATP production. In comparison, using an electron donor at a potential of 470 mV reduces the ΔG to −32 kJ/mol e^−^, lowering the theoretical yield to ~1.5 H^+^ exported per e^−^, and the larger ΔE^0'^ between the electrode and NAD(P)H requires at least four H^+^ translocated per NAD(P)^+^ reduced. This results in a balance of at least 80% and 20% for the electron utilization by the forward and reverse pathways. Thus, theoretically, at least twice as many electrons are needed to generate the same amount of reducing equivalents for CO_2_ fixation at the more positive potential. During growth in standard culture conditions, the acidophilic iron oxidizer *A. ferrooxidans* is predicted to have a ratio closer to 90/10% for the forward vs. reverse pathways, including the proton gradient necessary to generate ATP (30). Recent experimental evidence suggests that during growth on an electrode, the ratio of electron utilization by the forward to reverse pathways is close to 15:1 in *A. ferrooxidans* (10).

The majority of CBB cycle genes and pentose phosphate cycle genes appear to be more highly expressed at 310 mV. This likely reflects a higher rate of CO_2_ fixation at 310 mV compared to 470 mV. Furthermore, the higher relative expression of the Form IAc Rubisco genes and their associated carboxysomes at 470 mV may reflect the need to increase the efficiency of CO_2_ fixation due to the lower energy availability. However, even at 470 mV, the overall transcript abundance of Form IAq is still twice as high as Form IAc.

Among components possibly involved in ET at the outer membrane, the Cyc2 homolog encoded by Tel_03480 was more highly expressed at 470 mV in both metatranscriptomic and ddPCR samples, consistent with the lower energetic yield per electron. This protein has previously been implicated in EET in the acidophilic iron oxidizing bacterium *A. ferrooxidans* ATCC19859 (31–33). The E_M_ of Cyc2 in *A. ferrooxidans* is reported to be 560 mV at pH 4.8 (23). This is within the range that might be expected for a protein accepting electrons from a cathode at the potentials tested, although the midpoint potential of Cyc2 in “*Ca*. Tenderia electrophaga” is likely different. This strongly suggests that Cyc2 is involved in EET with the electrode in “*Ca*. Tenderia electrophaga”, although its exact role given the single heme and lack of transmembrane helices remains unknown.

Redundancy of electron transport chain components suggests large amount of metabolic flexibility in “*Ca*. Tenderia electrophaga” depending upon potentials of electron donors and local redox environment (Fig. 5). Thermodynamics dictates that the balance of electrons passing through the two branches of the ETC depends upon the electrode potential. One possible method of controlling this ratio would be modulating the abundance of the periplasmic links between EET and the ETC. Several proteins could potentially make up this link, but their roles are presently unclear. Hierarchical clustering of potential electron transfer components by expression leads to several distinct clusters (Fig. 3). One cluster contains the ACIII and Complex I genes, which suggests that these complexes may be functionally linked. This cluster also contains the Cyc1 homolog which suggests that it may facilitate the transfer of electrons to the “uphill” ETC, rather than to the terminal oxidase as proposed elsewhere (19, 30). The canonical cytochrome cbb_3_ oxidase is part of a cluster with succinate dehydrogenase (Complex II) and the cytochrome bc_1_ complex. This may indicate that they form part of a forward ETC, which would enable the use of stored reserves in the form of glycogen.

**Fig. 5.**
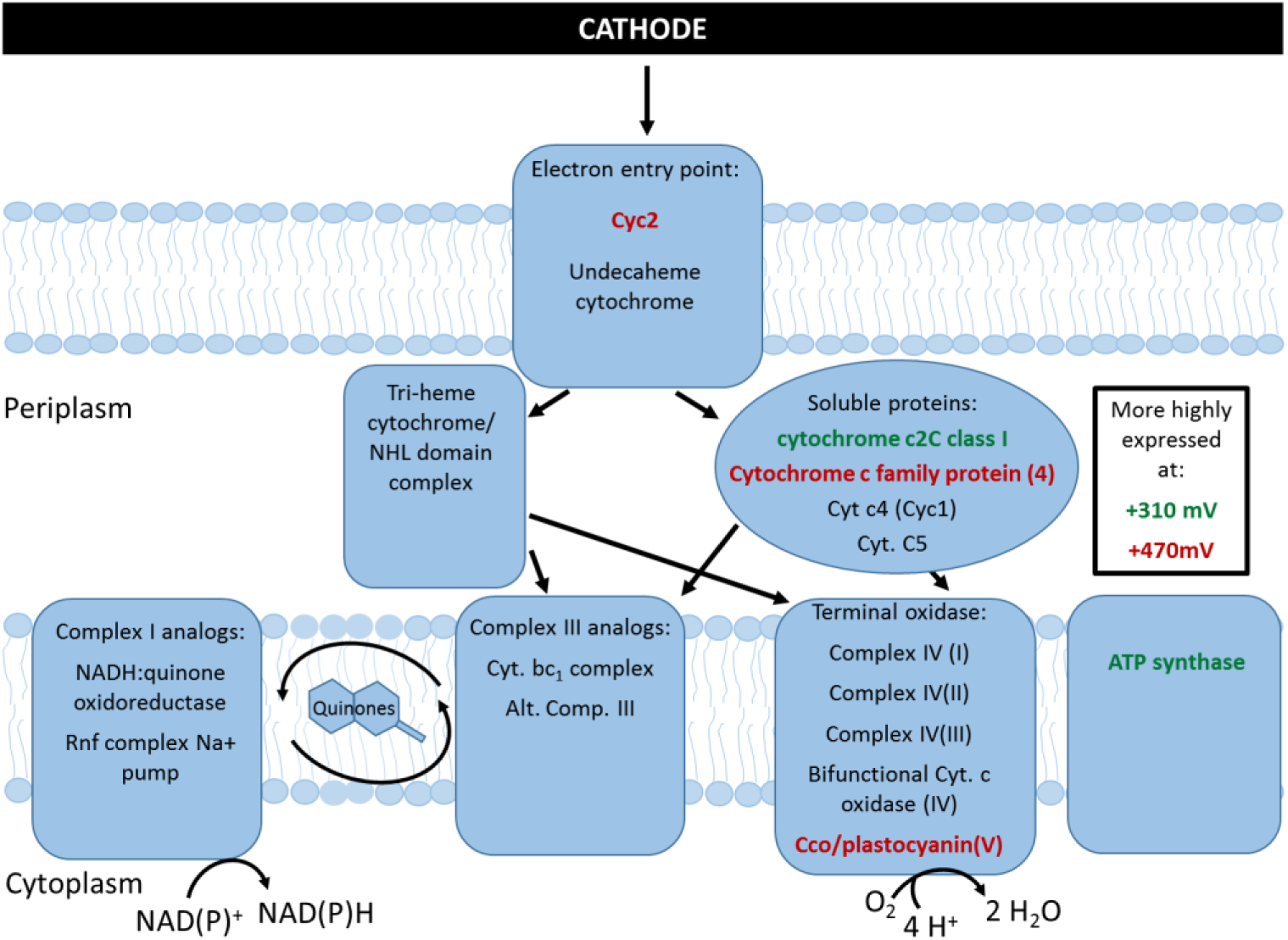
Schematic of potential EET/ETC routes in “*Ca*. Tenderia electrophaga”. Genes or complexes labeled in red are more highly expressed at 470 mV, those in green were more highly expressed at 310 mV.

Metatranscriptomics supports previous work by our group that identified “*Ca*. Tenderia electrophaga” as the electroautotroph. It appears to be the key member of the community and its predicted lifestyle as an electroautotroph is supported by a several key findings. “*Ca*. Tenderia electrophaga” was more active than any other organism and its activity was positively correlated with current density. Genes for proteins previously predicted to be involved in EET in “*Ca*.Tenderia electrophaga”, including Cyc2, were more highly expressed at 470 mV, when more electrons are needed to generate the same amount of PMF. The membrane bound ETC that is necessary to generate the ATP and reducing power for CO_2_ fixation was active at both potentials, but a potential complex IV analog and several that may be involved in the “downhill” branch of the ETC showed increased expression in response to changes in electrode potential. These changes were consistent with the need to use more electrons to obtain the same amount of energy. Most components of central carbon metabolism found were more highly expressed at 310 mV, suggesting a higher metabolic rate. The evidence for involvement of specific proteins or complexes in EET and ETC is correlative in nature, but narrows the list of potential targets for future confirmatory studies that are in progress. Further investigations with better temporal resolution may yield additional insights into the dynamics of the response to the changes in potential. Heterologous expression and biochemical measurements of predicted components of EET and the ETC may help identify functional roles. This work provides insight into the mechanisms for electroautotrophic growth in this biocathode community that will assist in engineering biocathode communities.

## METHODS

### Biofilm growth and sampling

Samples for RNA-seq were grown in artificial seawater medium (ASW) as previously described (8). Inocula consisted of cells detached from a 3 cm × 3 cm portion of a carbon cloth electrode and counted using an Accuri C6 flow cytometer (BD Bioscience, San Jose, CA) described previously (15). Sterile 2 L dual chambered microbial fuel cell reactors (Adams and Chittenden Scientific Glass) reactors containing a graphite coupon 3 cm × 9 cm × 0.2 cm were inoculated with standardized inocula estimated to contain 2 × 10^5^ cells.

Four reactors were inoculated with the same inoculum and grown under standard conditions (310 mV SHE), until they reached the maximum current (12 - 65 mA m^−2^). At this point, cyclic voltammetry (CV) was CV recorded from 610 mV to 260 mV and back to 610 mV at a scan rate of 0.2 mV/s. After CV, two reactors remained poised at 310 mV, and two were shifted to a suboptimal potential (470 mV SHE) where CV indicates that the rate of electron uptake is approximately half of that at the “optimal” potential. After 52 hours, the biofilm was scraped from ~3 × 9 cm portions of both sides of the cathode with a sterile razor blade, immediately immersed in RNA*later®* (life Technologies, Grand Island, NY) and stored at 4° C overnight prior to freezing at −20°C until total RNA extraction. This experiment was repeated 7 days later with a new inoculum. This gave two pairs of biological replicates for each treatment.

### RNA extraction, library prep, and sequencing

Biofilm RNA were extracted using an established protocol (34) with the following modification, instead of 0.8 g of 0.5 mm glass beads, 250 µL of Zirconia beads (Life Technologies) were used. The extracted RNA samples were subject to DNase treatment using Turbo DNase (Life Technologies) and cleaned up using RNA clean & concentrator^TM^-5 (Zymo Research). The DNase treatment was repeated once for each sample to ensure the removal of contaminating DNA. Ribosomal RNA was depleted from RNA samples using Ribo-Zero rRNA Removal Kit for Bacteria (Illumina, San Diego, CA). RNA library preparation was carried out using NEBNext^®^ Ultra RNA^TM^ Library Prep Kit for Illumina (New England Biolabs, Inc., Ipswich, MA) according to the manufacturer's recommended protocol. The sequence reads of RNA libraries were acquired on a MiSEQ^TM^ instrument under automated software control (version 2.2.0, Illumina, San Diego, CA). Raw reads that passed the initial quality filter were submitted to the NCBI Short Read Archive (http://www.ncbi.nlm.nih.gov/bioproject) under Bioproject PRJNA244670 experiment SRX978937 objects SAMN03462067 - SAMN03462074.

### Metatranscriptome alignment and statistical analysis of gene expression

Metatranscriptome reads were trimmed for quality using Trimmomatic 0.30 with a sliding window set to 5 bases with a minimum score of 25 and the first eight bases of the sequence and any leading bases with a quality score less than 20 were cropped (35). Reads shorter than 50 bases were discarded. To avoid inflated statistical confidence, only read one of the paired end reads was used, and if read one had been discarded during trimming, read two was used. Bowtie 2 (36) was used to align reads to the Biocathode-MCL metagenomic assembly available at http://www.ncbi.nlm.nih.gov under Bioproject PRJNA244670 (Malanoski et al. in prep), modified by removing contigs from the bins corresponding to PacBio sequenced genomes of “*Ca*. Tenderia electrophaga”, *Labrenzia* sp. CP4, and *Marinobacter* sp. CP1 (CP013099.1, CP013100.1, CP011929.1, CP011927.1, and CP011928.1) and contigs with >98% identical over >90% of their length. Single-end mode was used with the following options: -D 30 -R 3 -N 1 -L 15-i S,1,0.50 --score-min L,0,-0.15.

Read counts per contig were calculated using the idxstats function in samtools (37). Relative activity of each genome or metagenomic bin was calculated as the sum of RNA-seq reads mapping to the genome or bin divided by length of the genome or bin in nucleotides as a percent of activity in the library. Read counts per gene were calculated with HTseq-count. EdgeR (38, 39) was used to determine the statistical significance of differences in read counts for each annotated gene and pseudogene in “*Ca*. Tenderia electrophaga” with Benjamini-Hochberg adjustment to reduce the false discovery rate due to multiple hypothesis testing (40). A multifactor experimental design was used to separate changes in expression likely to be due to the different inocula from changes in expression resulting from the change in electrode potential (41). The sensitivity of this method to sequence coverage meant that a much smaller change in relative abundance of mRNAs was needed to reach the same FDR for genome bins with high coverage depth. Thus, the FDR used to determine a differentially expressed gene was FDR < 0.01. Values reported for gene expression at each condition are log_2_ of the number of reads per gene after normalization and variance stabilization transformation (42) (Table S4). Overall gene expression was calculated as EdgeR normalized gene expression minus the log_2_ of the length of the gene. For clustering genes by change in expression, gene expression was centered at 0 for each inoculum. Euclidean distance matrices were calculated, and genes were clustered using the Ward method implemented in the R package “pheatmap” version 0.7.7 (43).

Correlations between current density and relative abundance of mRNA from each metagenomic genome or bin were calculated as Pearson's product moment correlation coefficient (44). The critical value for a significant correlation for a two-tailed test with a sample size of n = 4 is 0.9 for a p-value of 0.1. A higher p-value cutoff for the correlation is justified because this is a different kind of statistical test and it assumes a linear relationship between the data, which is likely not the case for most genes. Furthermore, there is a large variability between samples of the same conditions (current magnitude, current increasing/decreasing, and community composition) that would prevent the near perfect correlation a p-value of 0.01 would require.

### Validation of differential expression with ddPCR

RNA samples from eight additional biological replicates were obtained from the same inoculum following the protocol above (four optimal and four suboptimal). After RNA extraction, a whole transcriptome amplification protocol was applied using the WTA-2 kit (Sigma) according to the manufactures instructions, performing a no template negative control. Individual ddPCR reactions for each primer set listed in Table S5 were set up using QX200 ddPCR EvaGreen Supermix (BioRad) and QX200 Droplet Generation Oil for EvaGreen (BioRad). Copy numbers of mRNAs for 19 genes were assessed as copy number per µl of reaction minus calculated copy number per µl of corresponding negative control reaction. Samples were normalized using a trimmed means approach to account for variation in the proportion of “*Ca*. Tenderia electrophaga” in the community, and significance of differential expression was assessed using two-tailed t-tests.

### Accession number(s)

All sequences produced in this study are available in the NCBI Short Read Archive under Bioproject number: PRJNA244670

## Acknowledgements

The authors would like to thank Drs. Nicholas J. Kotloski and Matthew D. Yates for helpful discussions about the processes and challenges unique to the electrode environment.

The opinions and assertions contained herein are those of the authors and are not to be construed as those of U.S. Navy, military service at large, or the U.S. government.

---

Table S1. Expression of ETC components in “Ca. Tenderia electrophaga”

Table S2. ddPCR results

Table S3. Expression of central carbon metabolism genes in “Ca. Tenderia electrophaga”

Table S4. Expression of all annotated “Ca. Tenderia electrophaga” genes

Table S5. Primer sequences for ddPCR

**Figure S1.**
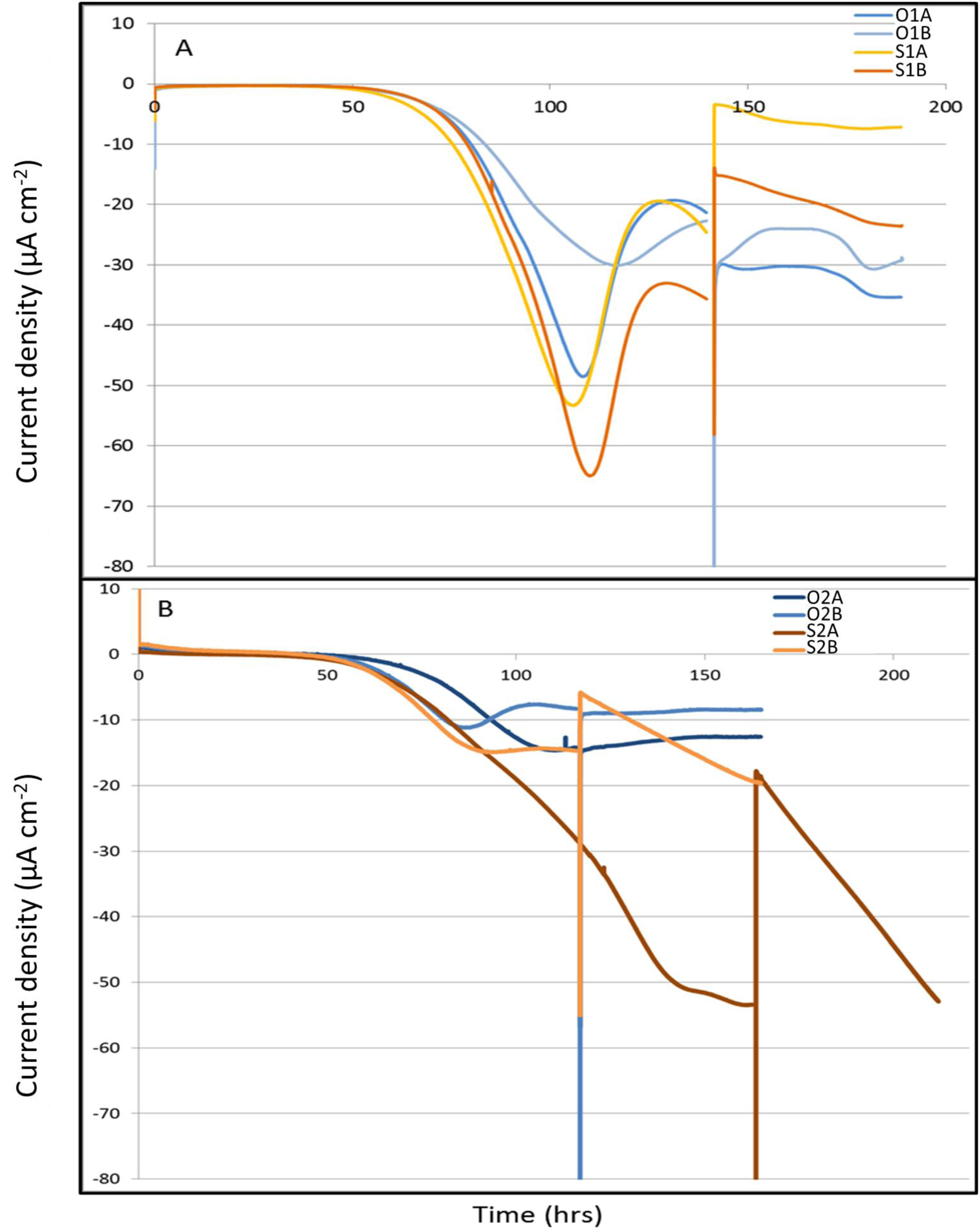
Chromoamperometry of biofilms grown at the 310 mv and either switched to 470 mV or continued at the optimal potential. Vertical lines indicate the time that the CV was measured and potential was switched. Sample ID string {O=optimal (310 mV), S=suboptimal (470 mV)}{Inoculum 1 or 2}{replicate A or B}.

**Fig. S2.**
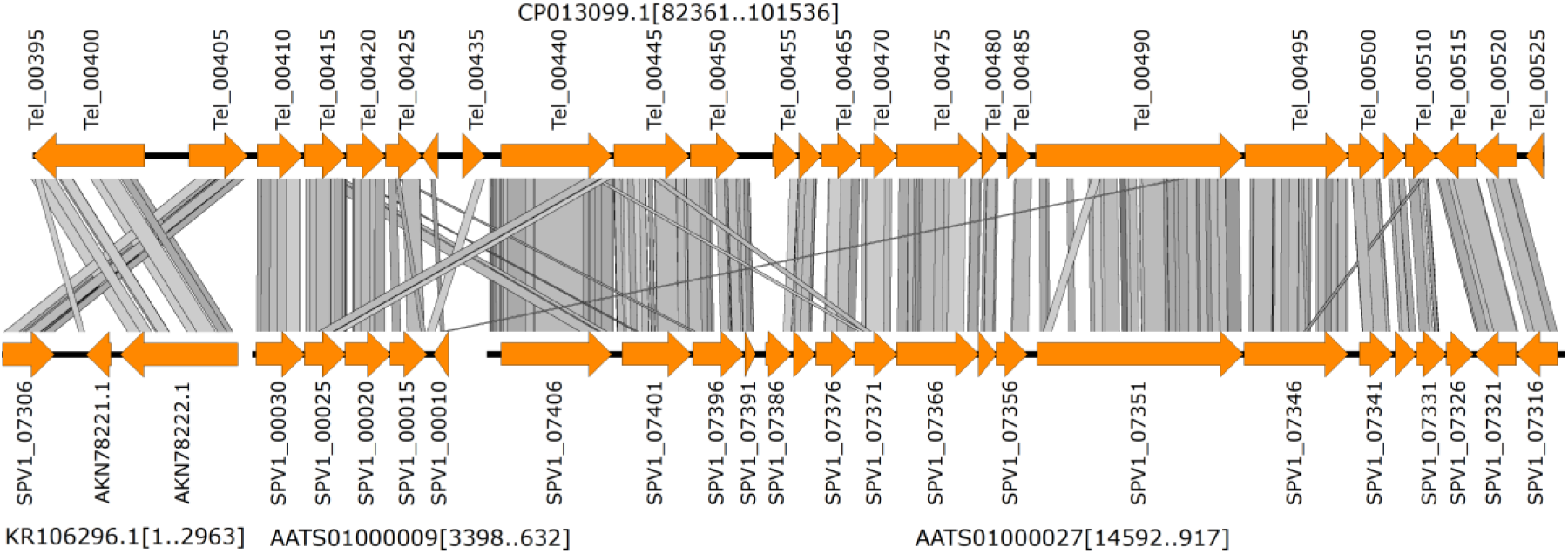
Synteny between “*Ca*. Tenderia electrophaga” and *Mariprofundus ferrooxydans* PV-1. The *M. ferrooxydans* genes are found on three contigs in the NCBI database, but two (KR106296.1 and AATS01000009) could be joined, as they have identical sequences at one end. Gray lines indicate regions of homology determined by a tBlastx search in EasyFig 2.2.2 with e-values of less than 1 × 10^−13^ with a minimum length of 30 bp (45). This region has been previously noted by Barco et al. to be syntenous within neutrophilic iron oxidizing bacteria (19).

